# The 2D Ising model, criticality and AIT

**DOI:** 10.1101/2021.10.21.465265

**Authors:** G. Ruffini, G. Deco

## Abstract

In this short note we study the 2D Ising model, a universal computational model which reflects phase transitions and critical phenomena, as a framework for establishing links between systems that exhibit criticality with the notions of complexity. This is motivated in the context of neuroscience applications stemming from algorithmic information theory (AIT). Starting with the original 2D Ising model, we show that — together with correlation length of the spin lattice, susceptibility to a uniform external field — the correlation time of the magnetization time series, the compression ratio of the spin lattice, the complexity of the magnetization time series — as derived from Lempel-Ziv-Welch compression—, and the rate of information transmission in the lattice, all reflect the effects of the phase transition, which results in spacetime pockets of uniform magnetization at all scales. We also show that in the Ising model the insertion of sparse long-range couplings has a direct effect on the critical temperature and other parameters. The addition of positive links extends the ordered regime to higher critical temperatures, while negative links have a stronger, disordering influence at the global scale. We discuss some implications for the study of long-range (e.g., ephaptic) interactions in the human brain and the effects of weak perturbations in neural dynamics.

## 1. Introduction

### 1.1. Network brain models

The human brain is a multiscale dynamical network. A body of evidence suggests that brain function emerges from interactions between specialized, spatially-segregated areas of large-scale networks. In such networks, nodes correspond to cortical or sub-cortical brain regions and edges correspond to either structural (i.e., direct connections) or functional (i.e., through synaptic or ephaptic interactions) couplings between these regions. Several computational studies have developed whole-brain network models to explore the relationship between brain function and its underlying connectivity [1, 2, 3, 4, 5, 6, 7, 8].

### 1.2. Critical phenomena and the 2D Ising model

More abstract frameworks from statistical physics can shed light into understanding emerging phenomena in large networks such as phase transitions in systems where nodes — neurons or cortical columns here — interchange information under the assumptions of the maximum entropy principle [9, 10, 11].

The description of systems with many degrees of freedom can be summarized by coarse-graining variables (describing macrostates), a step which introduces statistics into modeling. At so-called critical points, observable quantities such as diverging correlation length or susceptibility to external perturbations reveal singularities in the limit as the number of degrees of freedom goes to infinity. At these transitions, from order to disorder, there is a loss of sense of scale, with fractal properties in energy and information flow.

In this framework, elements such as neurons, columns or brain regions are modeled by spins (i.e., with two states, up or down, on or off) with nearest neighbor pair interactions, and the emerging statistical properties of large networks of these elements are studied under different conditions (temperature or excitability, or an external magnetic or electric field, see Figure 1). The prototypical simplest system in this context is the 2D Ising model, which features nearest neighbor interactions and a phase transition. This model has been shown to be universal, i.e., that all the physics of every classical spin model (with more general types of interactions) can be reproduced in the low-energy sector of certain “universal” models such as the 2D Ising model [12]. This fact reflects the intrinsic computational power of near-neighbor interactions. In fact, Ising models have been shown to be universally complete [13, 14], with a map between any given logic circuit to the ground states of some 2D Ising model Hamiltonian.

**Figure 1:**
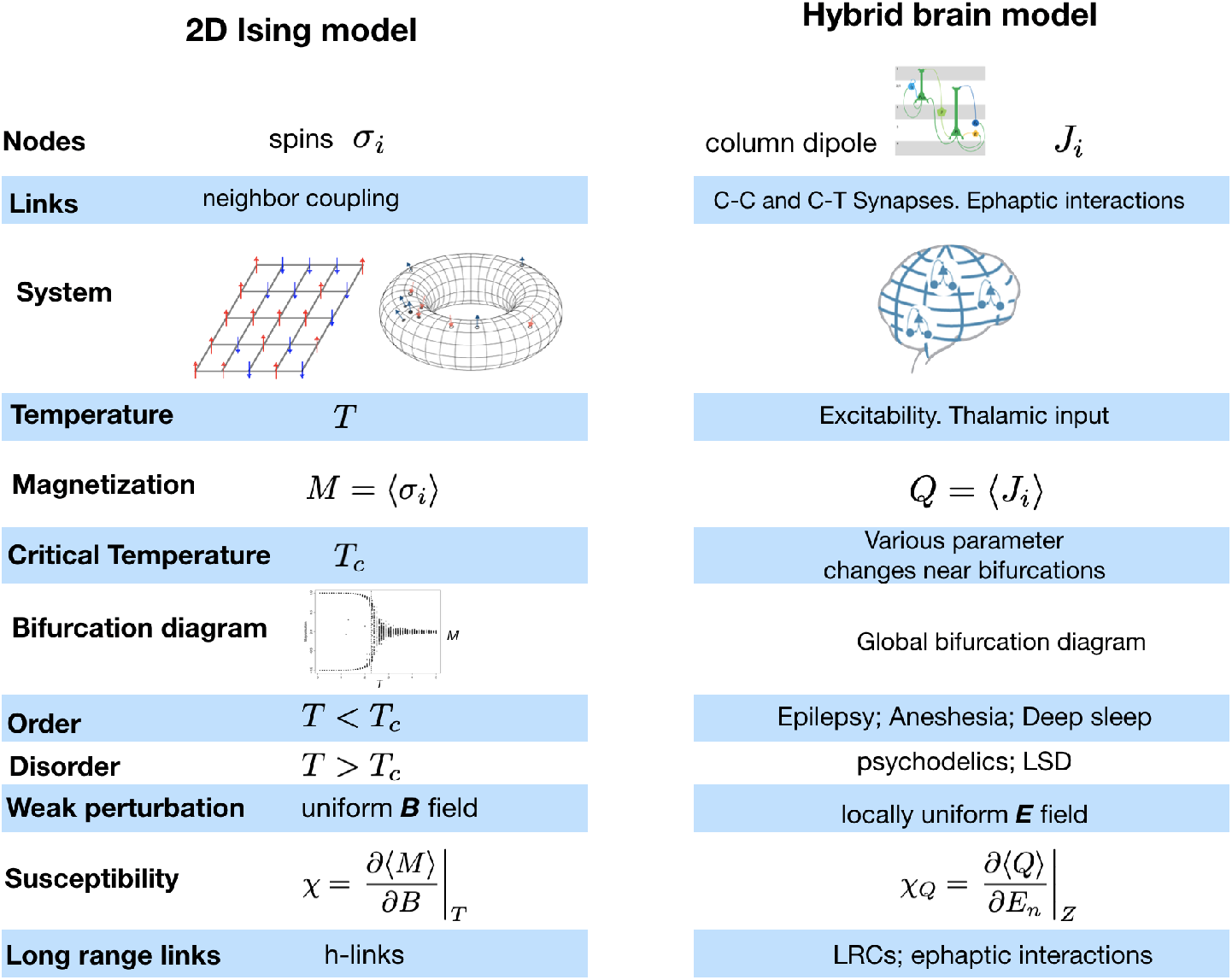
Parallels between the Ising 2D model and hybrid brain models (HMBs). LRC: long-range connectivity, synaptic or ephaptic.

On the other hand, the fact the brain exhibits characteristics of criticality that may be modeled by systems such as the Ising model is now well established, with ideas that go back to pioneers such as Turing, Bak [15] and Hopfield [16]. There is further evidence that the dynamics of the healthy brain occupy a sub-critical zone (see [17] and references therein).

While some recent studies have investigated the interpretation of fMRI data using the Ising formalism, our context is for its use with EEG data and for analysis/personalization of HBMs.

### 1.3. The amplification of weak perturbations by neuronal assemblies

Accumulating evidence from several decades of research suggests that endogenous or exogenous weak electric fields may influence neural processing, but how and where this may happen is not yet fully understood [18]. Weak electric fields such as the ones produced by non-invasive transcranial electrical current stimulation produce only very small perturbations of the transmembrane potential of the most sensitive neurons (elongated pyramidal cells). Understanding how these small (about 0.1–0.2 mV per V/m of electric applied [19] and significantly lower than the 20 mV depolarization required to bring a neuron from resting potential to spike threshold in vitro [20]), but relatively spatiotemporally homogenous single-cell effects collectively amplify is central to these questions. Two solutions can be envisioned.

Although membrane perturbations from weak fields are sub-threshold, nonlinear effects in coupled populations probably lead to an amplification of effects. For example, mathematical models have demonstrated the amplification of weak but coherent signals in networks of nonlinear oscillators (see, e.g., [21, 22, 23]) and, more specifically, in computational models of neural circuits [24, 25]). This effect is ultimately dependent on the coupling strength of network elements and their architecture, while noise can contribute to the enhancement of small but homogeneous perturbations in the network (array enhanced stochastic resonance). Thus, cooperative effects arising from noise and coupling in coupled systems can lead to an enhancement of the network response over that of a single element.

The second potential mechanism calls for the role of criticality, which is the one we directly study here. In fact, these two mechanisms, stochastic resonance and criticality, are probably closely related in the context of the human brain [26, 27].

### 1.4. Ephaptic interactions

When modeling systems as networks, the connectivity matrix of nodes plays a key role. Recently, the role of ephaptic interaction as an additional cortical coupling mechanism has been suggested. Ephaptic interactions refer to the direct effects on neurons of electric fields generated by the activity of other neurons [28]. Their main features are that they enable vert fast, bidirectional, propagation of information between cortical sites, influencing both local and synaptically distant regions as long as they are close in 3D space, and in a direction dictated by the state and orientation of the emitting and receiving populations (i.e., with effects that can be both excitatory and inhibitory). In [18], we studied the macroscopic electric field generated by cortical dipoles using realistic finite element modeling of the human brain. We found that modeled endogenous field magnitudes are comparable to those in measurements of weak but functionally relevant endogenous fields, or to those generated by transcranial current stimulation. The effects may thus play a role in linking synaptically distant regions, i.e., across sulci in the cortex.

### 1.5. Our aims here

As discussed above, a statistical network perspective appears of interest for two reasons. First, it may help understand the enhanced sensitivity of neuronal networks, which may find a direct analog in the increased susceptibility of Ising models near the critical temperature. Second, it may provide a tool to study the impact of long-range connections in the system (mimicking ephaptic effects) in properties such as critical temperature or susceptibility.

Here, we first consider the use of this model to study the links between the views afforded by statistical mechanics and algorithmic information theory, which is part of a unification program seeking to bring together algorithmic information theory, criticality and integrated information theory perspectives of the cognition (see Figure 2). We first aim to show that, at the abstract level, an Ising model can shed some light into how critical phenomena and complexity can be mathematically related, by studying algorithmic complexity of Ising model data near the critical temperature.

**Figure 2:**
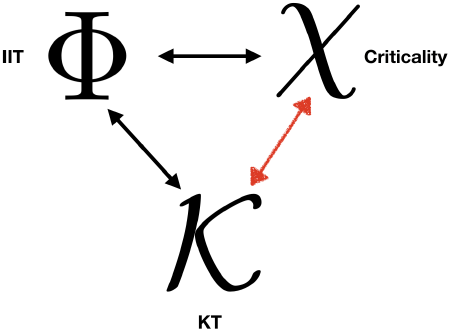
Algorithmic information or Kolmogorov unification program aiming to link algorithmic information theory, statistical mechanics and integrated information theory. Here we focus on the first two (red arrow).

Our second task is to then investigate the impact of short and long range “ephaptic” links on the classical 2D Ising model with local couplings (the original model) on phenomena such as phase transitions and complexity. The impact on susceptibility to weak perturbations will also be studied. As discussed, this is of interest due to the presence of such long range interactions in the human brain, both structural or electrical in nature (ephaptic interactions).

In the next sections, we first describe the model used and the metrics we will employ to study its dynamics. We then provide the results of simulations and discuss the results.

## 2. Materials and Methods

We first describe the original 2D Ising model with fields.

### 2.1. The 2D Ising model

The Ising model is given by the energy or Hamiltonian of the 2D lattice of side length *L, S* = {*σ*_*i*_|*i* = 1,.., *N*_*l*_ ≡ *L*^2^},

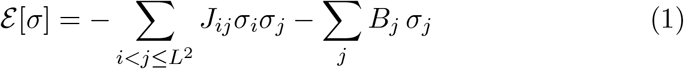

where *σ*_*i*_ denotes the orientation of each spin in the lattice (±1), *J*_*ij*_ is the coupling matrix (reflecting nearest neighbor coupling only with pairs counted once) and *B* is an external magnetic field (see Figure 3) that may be inhomogeneous.

**Figure 3:**
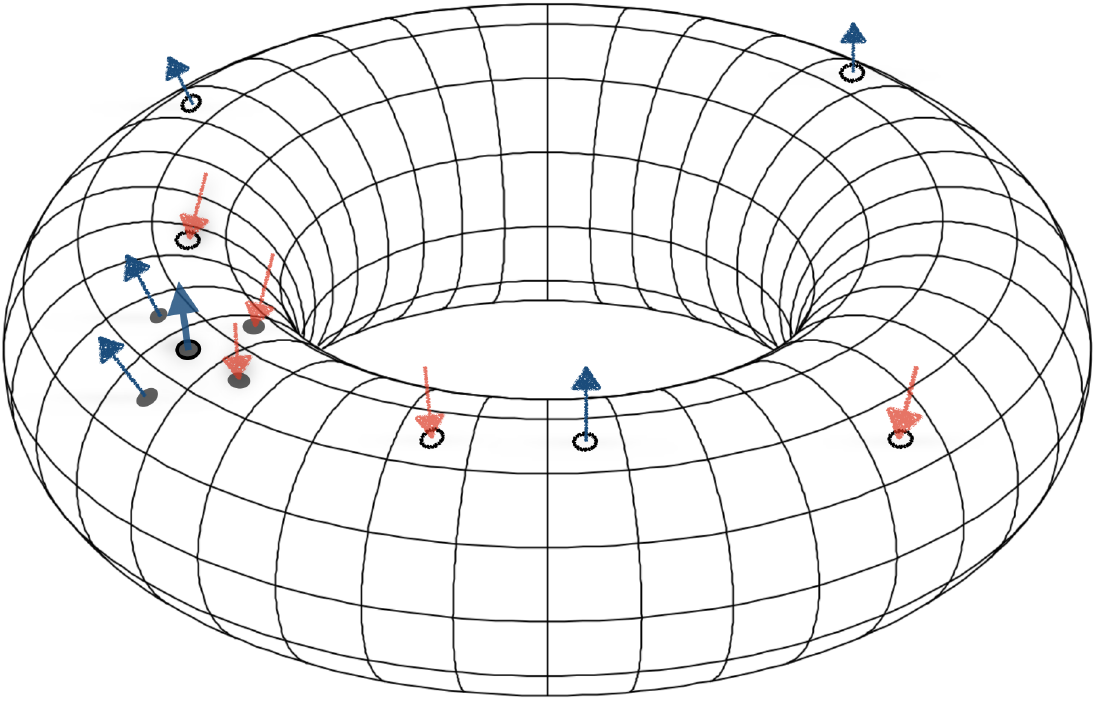
Ising model on 2D torus geometry. The torus has actually the same number of sites in each of its to directions — the lattice is square with periodic boundary conditions on the *J*_*ij*_. Nearest neighbors and remote spins that may be linked are depicted. A “focus” spin is highlighted on the left (solid circle with border) with its four nearest neighbors. Distant spins to which it may couple are also shown.

We can rewrite this using 2d indices as (we assume a uniform *B* field for simplicity)

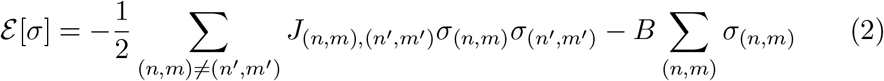

The energy of a single spin is

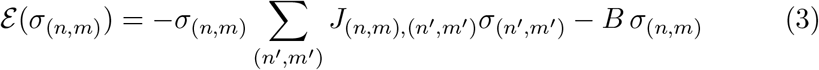

### 2.2. Hyperlinks (h-links)

In order to study the impact of additional sparse short or long-range couplings, we define a linking probability and create additional elements in the *J*_*ij*_ matrix. Holding the number of additional links fixed, we then study three different scenarios: a) uniformly selected random links, b) long-range links on the “antipodal” points in the toroidal manifold, and c) near-range links. The probability of adding a new link to a given site used was of 0.01%.

### 2.3. Implementation

The model has been implemented in Python 3 with the Numpy [29] and Scipy [30] libraries. Starting from a random lattice configuration, the system is evolved using a modified Metropolis algorithm where we update the state of a quarter of the spins (randomly chosen) at each time step. The number of time steps used is of ∼1–2 ×10^5^.

### 2.4. Observables

The main global observable is the lattice average magnetization *M* over the spin lattice *S*,

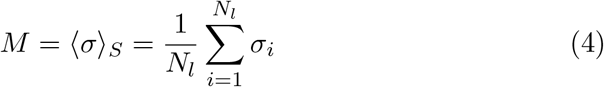

and lattice energy, where recall *N*_*l*_ is the number of points in the lattice. This is a function of time as the lattice evolves. Relying on ergodicity, we can compute time averages instead of ensemble averages.

#### 2.4.1. Autocorrelation length of lattice and time

Autocorrelation length or time are computed from data by first generating the autocorrelation function, and then computing from it the integrated autocorrelated time or length, e.g., 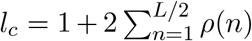 [31, 32]. The auto-correlation function of the lattice is computed along the two main axes and averaged.

#### 2.4.2. Compression ratio of magnetization time series or lattice data

The compression ratio is computed essentially as described in [33], i.e., compressing the quantized time series using the Lepel-Ziv-Welch algorithm and computing the ratio of compressed string length vs original string length. For (binary) lattice data, the array is flattened first. For time series data, quantification is done in 10 levels, (i.e., with digits 0-9).

As discussed in [33], two variants of this metric are computed, *ρ*_0_ and *ρ*_1_ = *ρ*_0_ − *H*_0_, with *H*_0_ the first order entropy. The second metric provides a measure of second and higher order effects on entropy rate, which can be interpreted as the extra apparent extra entropy (bits/char) incurred by using first order methods instead estimating the true entropy rate.

### 2.5. Magnetic susceptibility and heat capacity

These are computed from statistical mechanics considerations. Let

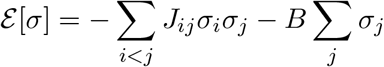

and

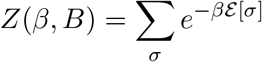

be the partition function. The average magnetization is the ensemble average of lattice magnetization,

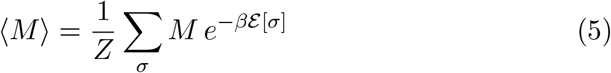

Then we can express (see, e.g., [34])

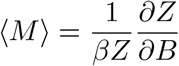

and following similar considerations

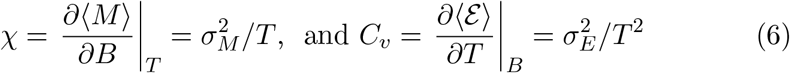

where the standard deviation computation is taken over the ensemble (i.e., time averages here).

### 2.6. Susceptibility to spin perturbations

We have also studied the impact of spin flips on lattice dynamics as a function of temperature. In order to do this we clamp the time series at a spin location to a randomly generated time series, and we study the correlation of this time series with that of a spin at a nearby location. Figure 7 displays the correlation and its p-value for different temperatures, highlight the increase in information transmission at the critical temperature.

### 2.7. Information transmission

Information transmission in Ising models of the brain has been studied before [35], finding that it is maximal at the critical temperature. Here we will study information transmission in the classical Ising model with h-links. Our approach is simple: we inject information at a spin site (clamping it down) and study the correlation of its forced activity at a displaced site, looking for maximal correlation at some delay Δ*τ*.

## 3. Results and discussion

### 3.1. Impact of positive hyperlinks

In Figure 4 the results from a 40×40 lattice are displayed, with and without long range hyper links. The first conclusion is that adding a few positive-coupling hyperlinks increases the critical temperature and extends the sub-critical range, as reflected in the magnetization, heat capacity and susceptibility. The effect is larger for larger lattices given a fixed percent of new links (0.01% here).

**Figure 4:**
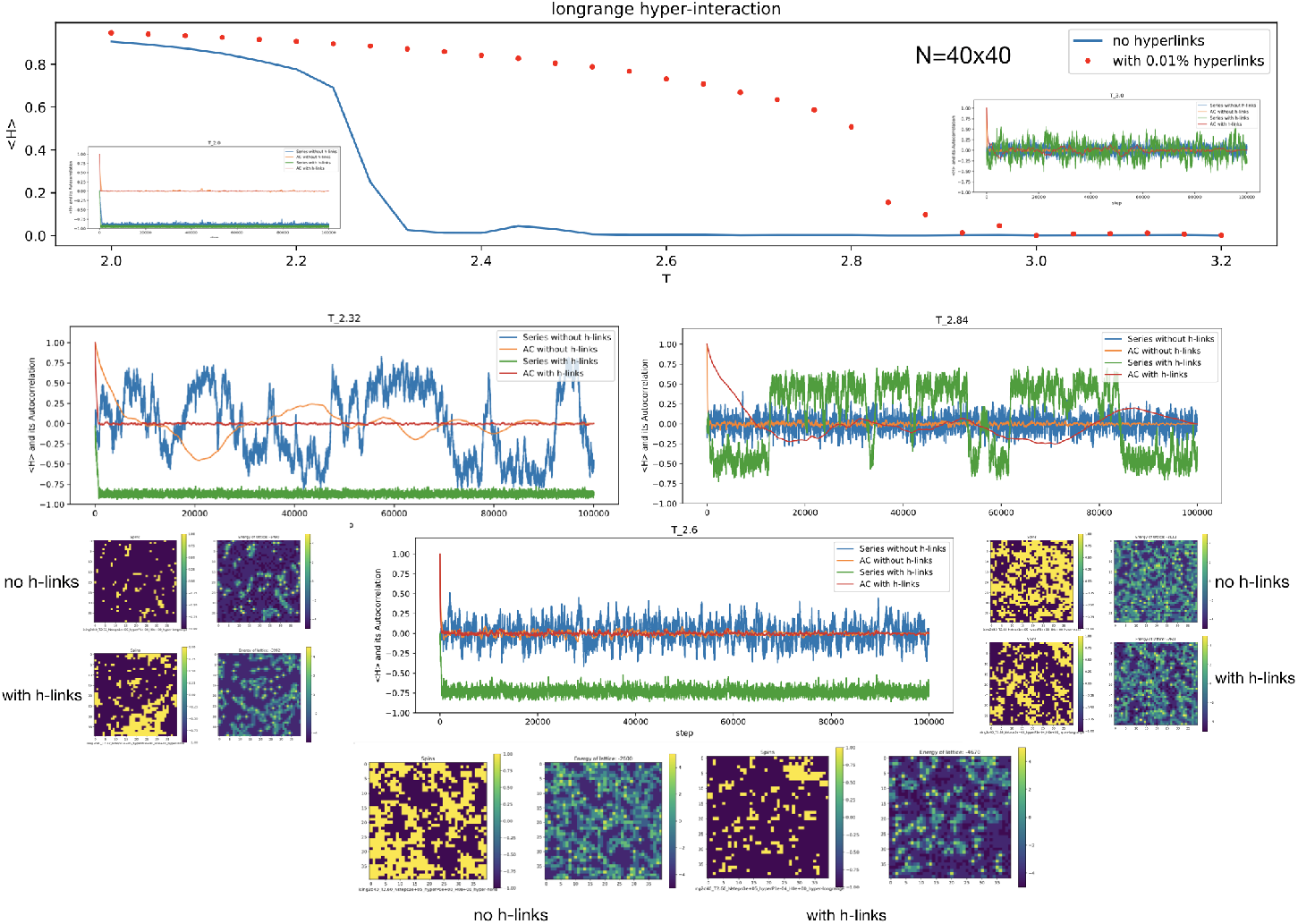
Results from simulation: magnetization as a function of temperature in a 40×40 lattice with and without long-range, sparse (0.01% probability of coupling) positive couplings on the critical temperature.

The effect of the hyperlinks is also seen on the complexity metrics. Figure 5 displays the compression ratio and autocorrelation length and time of lattice and magnetization time series. The addition of hyperlinks creates a local minimum in the compression ratio of the time series.

**Figure 5:**
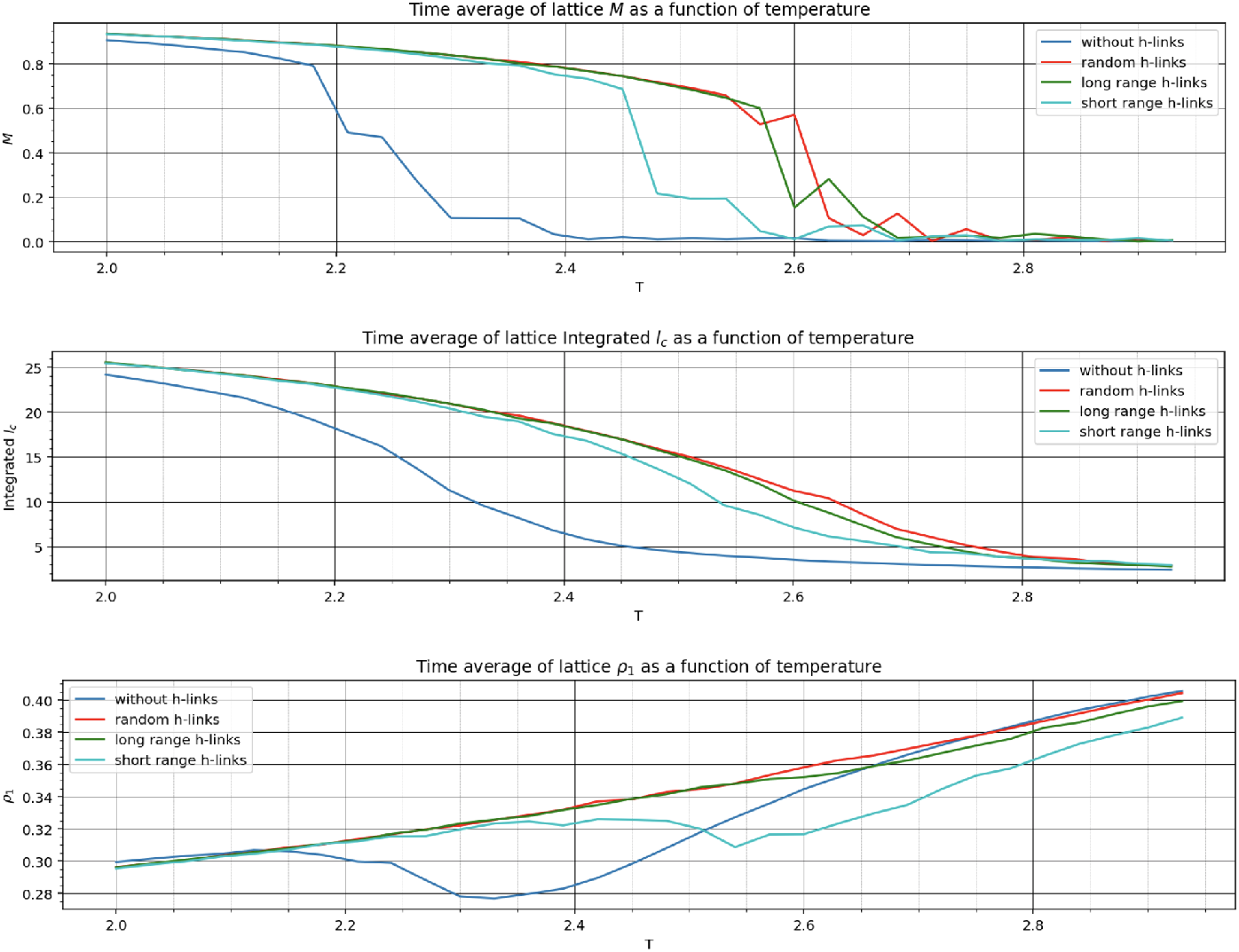
Results from simulation: lattice magnetization, correlation length and compression ratio as a function of temperature (30×30 lattice, 1e5 steps) with and without positive links.

The addition of a new links amounts, in some sense, to a change in the topology of the lattice, i.e., increasing its genus. However, the analogy is not perfect, as in a proper topological sense, we should reconfigure more elements of the connectivity matrix to ensure each node has the same number of nearest neighbors (4).

### 3.2. Impact of negative hyperlinks

The addition of negative coupling h-links has a very strong effect, and effectively reduces the phase transition temperature point.

### 3.3. Complexity and critical temperature

The results of running a 30×30 lattice model can be seen in Figure 5 and Figure 6. The critical temperature of the classical model does not appear to be affected by lattice size, but the addition of extra links has a strong impact, in particular of long range links.

**Figure 6:**
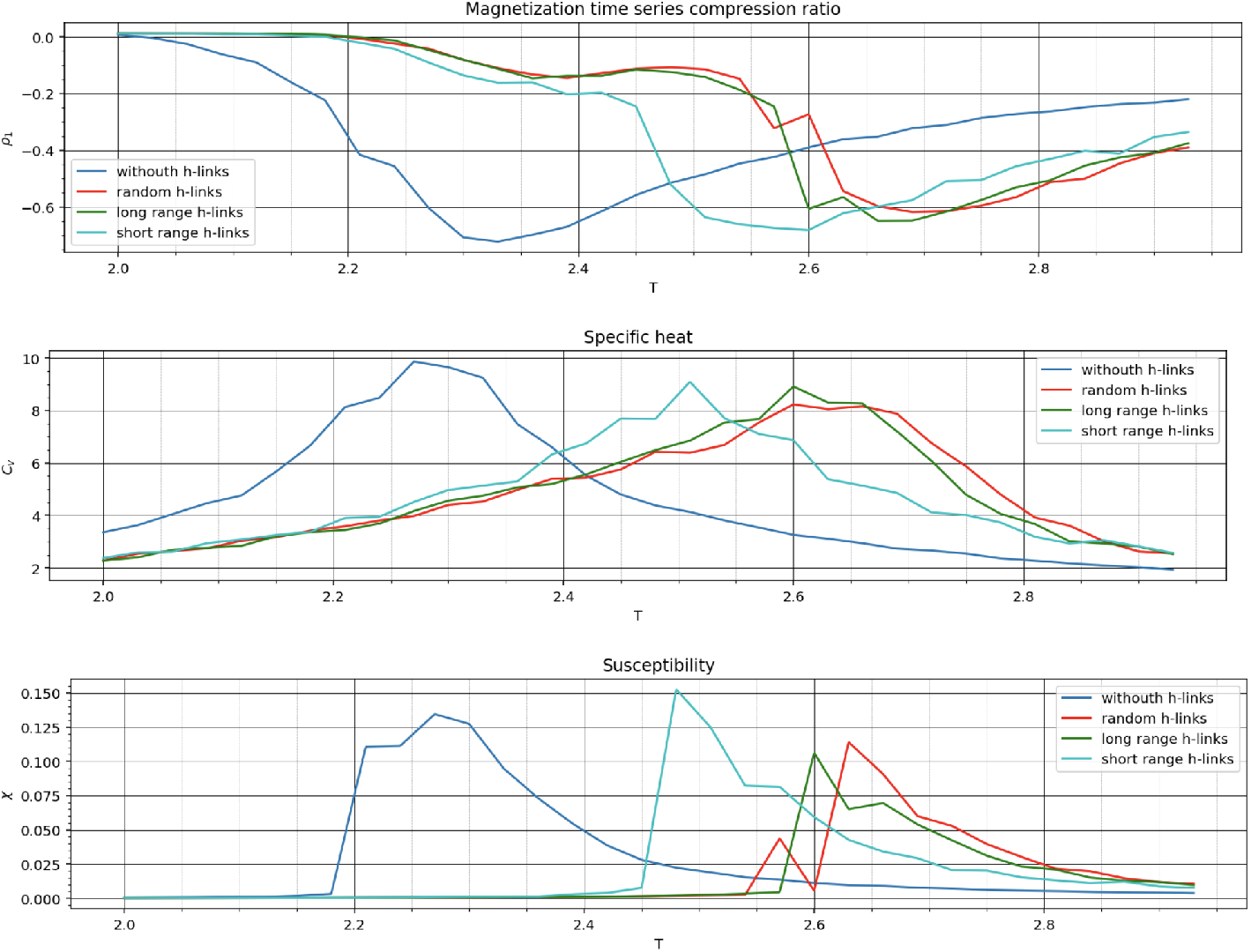
Results from simulation: magnetization time series compression ratio, specific heat and susceptibility as a function of temperature (30×30 lattice, 1e5 steps).

**Figure 7:**
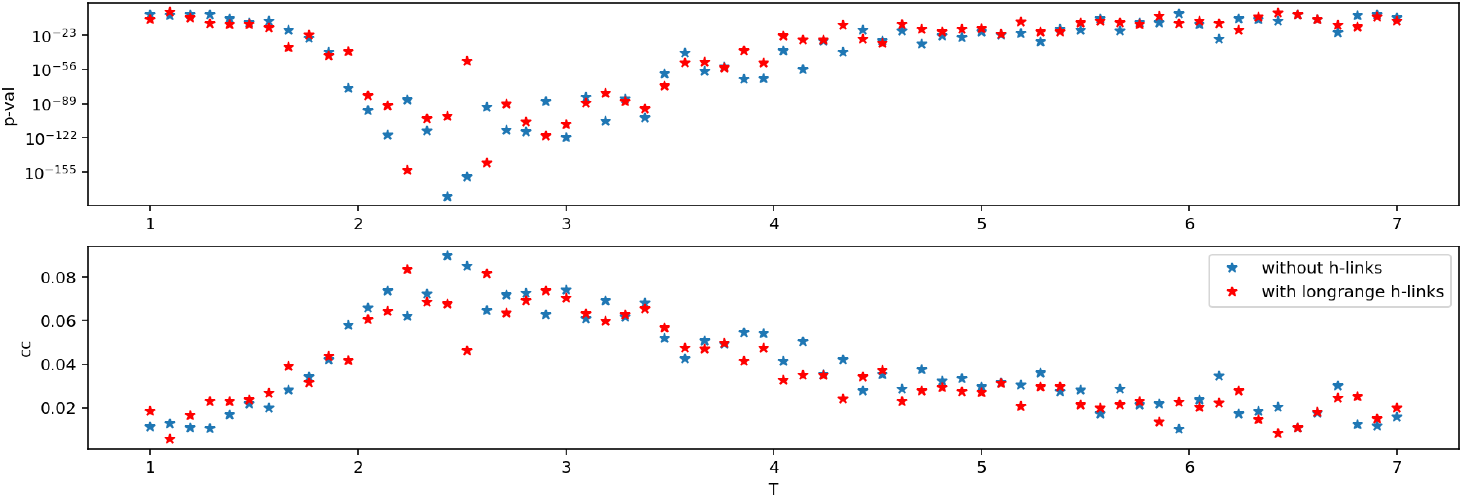
Information transmission as a function of temperature measured by p-value of correlation of the injected noise at a site with activity at another site displaced two steps. (30×30 lattice, 1e5 steps). Top: p-value. Bottom: cross-correlation coefficient.

**Figure 8:**
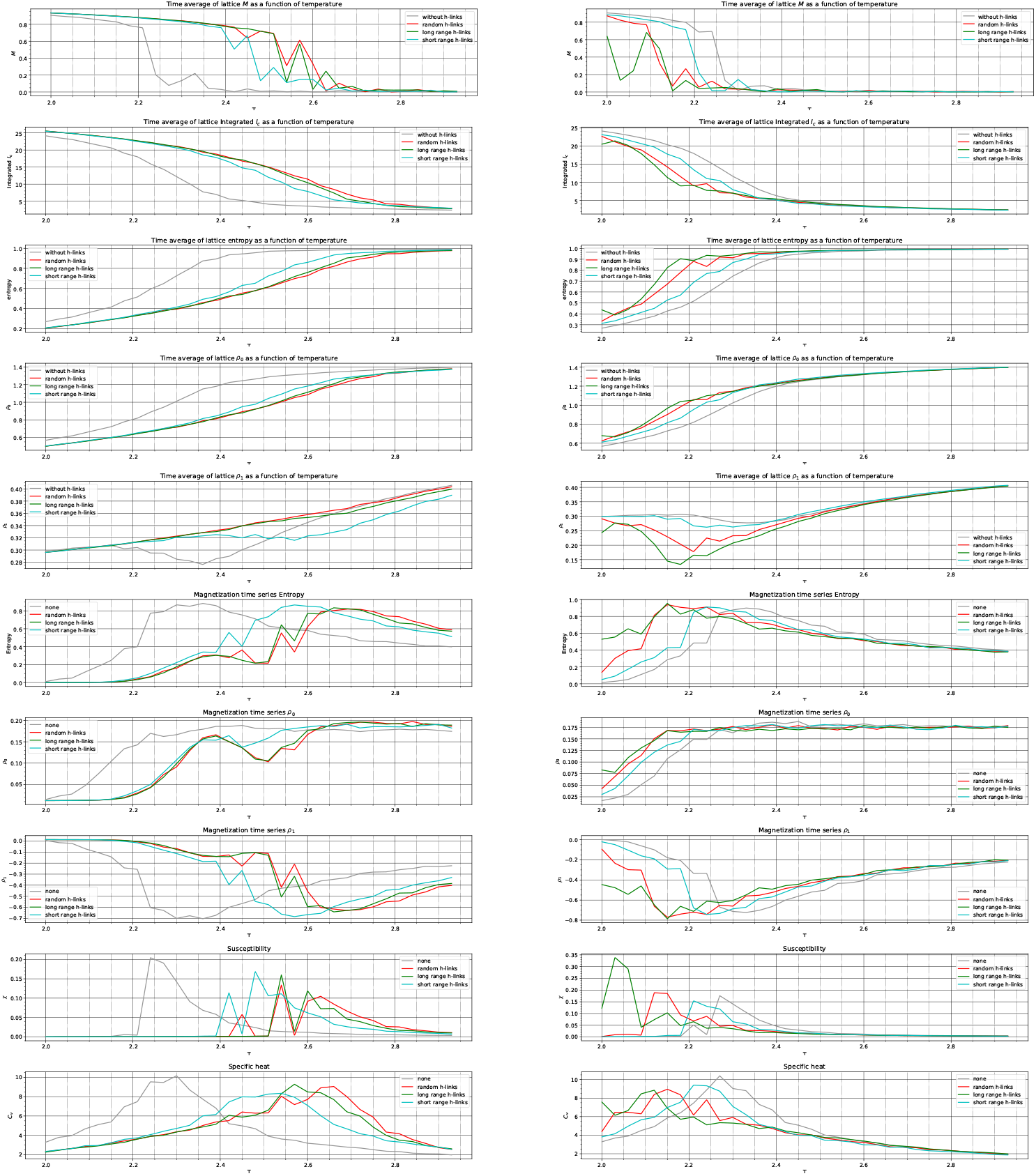
Results from simulation—lattice and time series metrics. Left: positive (p=1e-4) couplings. Right: negative couplings (p=2e-5). (30×30 lattice, 1e5 steps).

### 3.4. Information transmission is maximal at the critical temperature unaffected by long range h-links

Figure 7 displays the correlation statistics as a function of temperature. Information transmission is maximal near the critical temperature of the standard model, irrespective of the presence of h-links.

## 4. Conclusions

The main conclusion from this study is that the Ising model is very sensitive to the presence of a few long range (positive) connections. This has substantial impact on the critical point of the system. Negative connections seem to have an even stronger effect.

The second conclusion is that the transition from order to disorder defined by the critical boundary has a fingerprint on the entropy of the dynamics of system as measured by Lempel Ziv complexity, an effect that can be seen in the entropy of variables such as the magnetization time series.

Finally, we have looked at information transmission in the lattice as function of temperature, verifying that it peaks at the critical temperature, as expected. The presence of long range links does not seem to have an effect on the temperature at which the peak is found.

### 4.1. Relevance to hybrid brain modeling

One of the key question in HBM personalization is how to fix model parameters. An interesting line of work can be foreseen using Ising models derived from either cortically mapped EEG data, or from the fitted model. From a fitted HBM we can derive an Ising model and study it from the point of view of criticality for further insights into how to intervene. For example, we can binarize the time series associated to each NMM dipole field *J* (*t*) and then fit a Hamiltonian to reproduce the energy landscape in the data as in [36], and to see how far from the critical temperature it is.

It is possible that a necessary condition for a healthy brain will be to operate near but under its critical temperature. It has been argued that some brain states (e.g., psychodelics) or pathologies (fibromyalgia) can be attributed to *critico-patologies* — being too close [37, 17, 38] or below the optimal zone. Several neurodenerative diseases have been characterized as displaying less complexity (e.g., AD or PD, see e.g., [39] and references therein).

In future research we plan to use personalized Ising models to study both energy landscapes and associated temperature to see how close to criticality the system is.

### 4.2. A personalized critical temperature for the individual brain

Ising models have been used to model brain activity (see., e.g., [36]). Here we would suggest to proceed as follows. Use a parcellation of the cortex and create an associated 2D Ising (classical) model, but with connectivity defined by the connectome (see, e.g., [40]). Add h-links as estimated by the ephaptic coupling index (see [41]) or related considerations. Compute the critical temperature and modified susceptibility of the created Ising model to estimate the effect of perturbations by external fields (globally or locally defined by, e.g., realistic electric field modeling). This can be done at the global level, in each parcel.

### 4.3. Ising and individual or brain state dependence to tES

As we have seen, criticality may be an important concept in tES. There is a hypothesis we can explore related to the inter-individual variability in tES responses, or something called “brain state dependence” of stimulation effects. Could this be related to the *T*_*c*_ of each patient? We can fit an Ising model to each person and estimate distance to critical temperature. There are different routes to do this, and different data sources we can use. A fitted HBM can be the starting point. Or we can create an Ising model directly from data (EEG or fMRI or other as in [36]), and include or not DTI information. The hypothesis would be that what are traditionally called “responders” in brain stimulation (tES or TMS) are those that happen to be near *T*_*c*_ when stimulation is applied.

In order to account for the effect of the electric field, we need to adapt it to the Ising modeling framework. The effect of an electric field can be modeled as a shift of the firing rate of neuronal populations. If we think of the spin as a discretized version of the local cortical dipole, we can connect the two worlds, for example, by selecting a bandpass for the signals (fMRI, EEG, MEG or others) and discretize the power envelope to reflect an up or down state (−1 being a state of low activity).

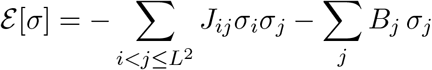

with

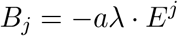

where *E*^*j*^ is the vector E field value at node *j* in the physical mesh.

## Acknowledgements

This project has received funding from the European Research Council (ERC) (Galvani) under the European Union’s Horizon 2020 research and innovation programme (grant agreement No 855109) and from the European Union’s Horizon 2020 research and innovation programme under grant agreement No 101017716 (Neurotwin).

